# RNA polymerase errors cause splicing defects and can be regulated by differential expression of RNA polymerase subunits

**DOI:** 10.1101/026799

**Authors:** Lucas B. Carey

## Abstract

Errors during transcription may play an important role in determining cellular phenotypes: the RNA polymerase error rate is >4 orders of magnitude higher than that of DNA polymerase and errors are amplified >1000-fold due to translation. However, current methods to measure RNA polymerase fidelity are low-throughout, technically challenging, and organism specific. Here I show that changes in RNA polymerase fidelity can be measured using standard RNA sequencing protocols. I find that RNA polymerase is error-prone, and these errors can result in splicing defects. Furthermore, I find that differential expression of RNA polymerase subunits causes changes in RNA polymerase fidelity, and that coding sequences may have evolved to minimize the effect of these errors. These results suggest that errors caused by RNA polymerase may be a major source of stochastic variability at the level of single cells.

The information that determines protein sequence is stored in the genome but that information must be transcribed by RNA polymerase and translated by the ribosome before reaching its final form. DNA polymerase error rates have been well characterized in a variety of species and environmental conditions, and are low, on the order of one mutation per 10^8^ - 10^10^ bases per generation^1-3^. In contrast, RNA polymerase errors are uniquely positioned to generate phenotypic diversity. Error rates are high (10^−6^-10^−5^)^4–7^, and each mRNA molecule is translated into 2,000 – 4,000 molecules of protein^8,9^, resulting in amplification of any errors. Likewise, because many RNAs are present in less than one molecule per cell in microbes^10,11^ and embryonic stem cells^12^, an RNA with an error may be the only RNA for that gene; all newly translated protein will contain this error. Despite the fact that transient errors can result in altered phenotypes^13,14^, the genetics and environmental factors that affect RNA polymerase fidelity are poorly understood. This is because current methods for measuring polymerase fidelity are technically challenging^4^, require specialized organism-specific genetic constructs^15^, and can only measure error rates at specific loci^16^.

To overcome these obstacles I developed MORPhEUS (Measurement Of RNA Polymerase Errors Using Sequencing), which enables measurement of differential RNA polymerase fidelity using existing RNA-seq data **(Figure 1).** The input is a set of RNA-seq fastq files and a reference genome, and the output is the error rate at each position in the genome. I find that RNA polymerase errors result in intron retention and that cellular mRNA quality control may reduce the effective RNA polymerase error rate. Moreover, our analyses suggest that the expression level of the RPB9 Pol II subunit determines RNA polymerase fidelity *in-vivo.* Because it can be run on any existing RNA-seq data, MORPhEUS enables the exploration of a previously unexplored source of biological diversity in microbes and mammals.

**Figure 1.**
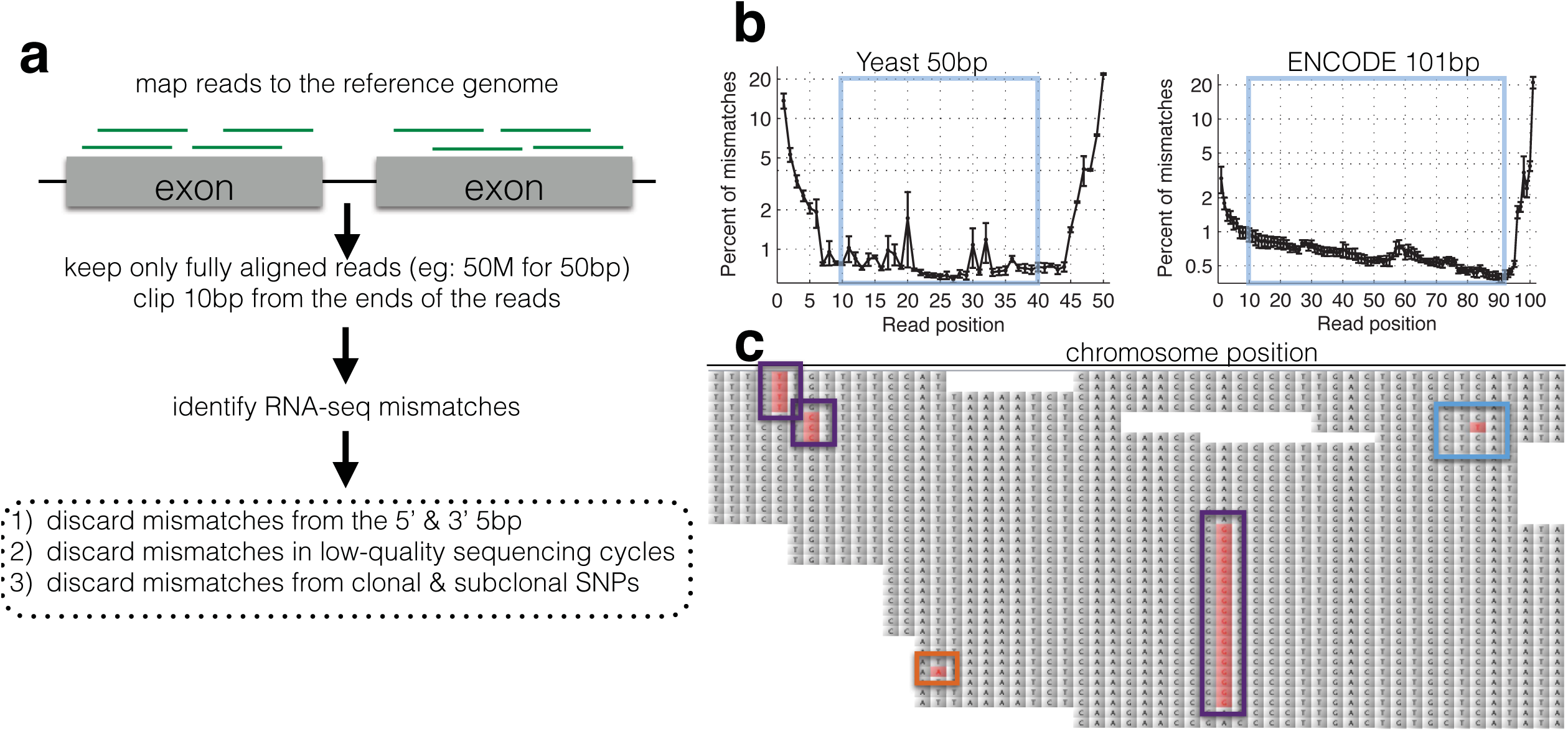
A computational framework to measure relative changes in RNA polymerase fidelity. **(a)** Pipeline to identify potential RNA polymerase errors in RNA-seq data. High quality full-length RNA-seq reads are mapped to the reference genome or transcriptome using bwa, and only reads that map completely with two or fewer mismatches are kept. **(b)** Then 10bp from the front and 10bp from the end of the read are discarded as these regions have high error rates and are prone to poor quality local alignments. **(c)** Errors that occur multiple times (purple boxes) are discarded, as these are likely due to sub-clonal DNA mutations or sequences that sequence poorly on the HiSeq. Unique errors in the middle of reads (cyan box) are kept and counted.

Technical errors from reverse transcription and sequencing, and biological errors from RNA polymerase look identical (single-nucleotide differences from the reference genome). Therefore, a major challenge in identifying SNPs and in measuring changes in polymerase fidelity is the reduction of technical errors^17–19^**(Figure 1)**. First, I map full length (untrimmed) reads to the genome, and discard reads with indels, more than two mismatches, reads that map to multiple locations in the genome, and reads that do not map end-to-end along the full length of the read. I next trim the ends of the mapped reads, as alignments are of lower quality along the ends, and the mismatch rate is higher, especially at splice junctions. I also discard any cycles within the run with abnormally high error rates, and bases with low Illumina quality scores **(Figure 1 – figure supplement 1)**. Finally, using the remaining bases, I count the number of matches and mismatches to the reference genome at each position in the genome. I discard positions with identical mismatches that are present more than once, as these are likely due to subclonal DNA polymorphisms or sequences that Illumina miscalls in a systematic manner^20^ **(Figure 1 - figure supplement 2)**. The result is a set of mismatches, many of which are technical errors, some of which are RNA polymerase errors. In order to determine if RNA-seq mismatches are due to RNA polymerase errors it is necessary to identify sequence locations in which RNA polymerase errors are expected to have a measurable effect, or situations in which RNA polymerase fidelity is expected to vary.

I reasoned that RNA polymerase errors that alter positions necessary for splicing should result in intron retention, while sequencing errors should not affect the final structure of the mRNA **(Figure 2a)**. However, mutations in the donor and acceptor splice sites also result in decreased expression^36^, and therefore are difficult to measure using RNA-seq. I therefore used chromatin-associated and nuclear RNA from Hela and Huh7 cells^37^, and extracted all reads that span an exon-intron junction for introns with canonical GT and AG splice sites, and measured the RNA-seq mismatch rate at each position. I find that errors at the G and U in the 5’ donor site, and at the A in the acceptor site are significantly enriched relative to errors at other positions **(Figure 2b)**, and to errors trinucleotides present in the splicing motifs in the human genome **(Figure 2 - figure supplement 1)** suggesting that RNA polymerase mismatches can result in changes in transcript isoforms. The ability of RNA polymerase errors to significantly affect splicing has been proposed^22^ but never previously measured.

**Figure 2.**
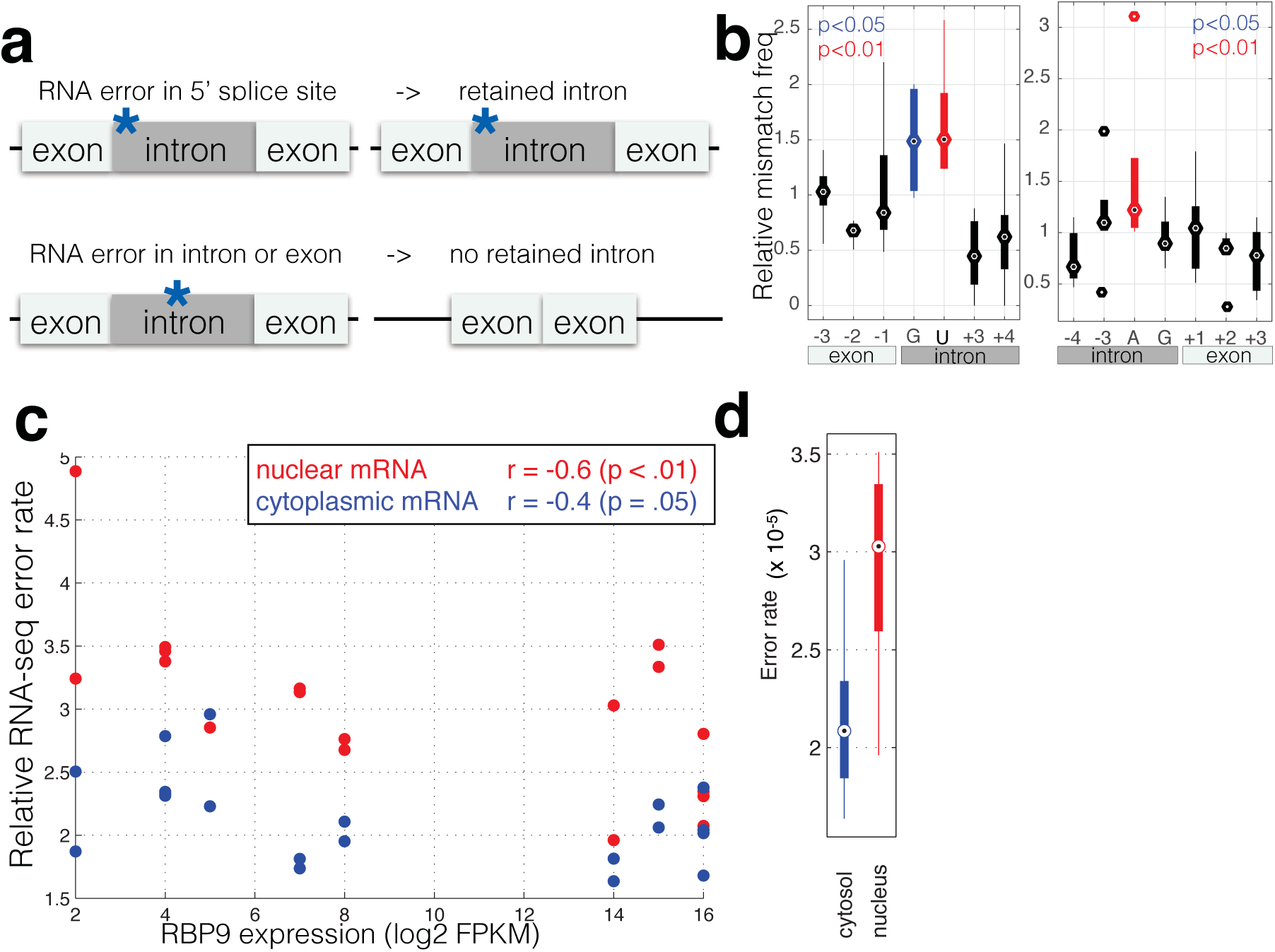
RNA polymerase errors cause intron retention and error rates are correlated with RPB9 expression. **(a)** RNA polymerse errors at the splice junction should result in intron retention, as DNA mutations at the 5’ donor site are known to cause intron retention. **(b)** Shown are the RNA-seq mismatch rates at each position relative to the 5’ donor splice site, for sequencing reads that span an exon-intron junction. Mismatch rates from chromatin associated and nuclear RNAs are higher at the 5’ and 3’ splice sites, suggesting that RNA polymerase errors at this site result in intron retention. **(c)** For all ENCODE cell lines, RPB9 expression was determined from whole-cell RNA-seq data, and the RNA-seq error rate was measured separately for the cytoplasmic and nuclear fractions. **(d)** The RNA-seq error rate is higher (paired t-test, p=0.0019) in the nuclear than the cytoplasmic fraction, suggesting that quality control mechanism may block nuclear export of low quality mRNAs.

RPB9 is known to be involved in RNA polymerase fidelity *in vitro* and *in vivo*^15,23^. I therefore reasoned that cell lines expressing low levels of RPB9 would have higher RNA polymerase error rates. Consistent with this, I find that RPB9 expression varies 8-fold across the ENCODE cell lines, and this expression variation is correlated with the RNA-seq error rate **(Figure 2c, Figure 2 - figure supplement 2)**. This suggests that low RPB9 expression may cause decreased polymerase fidelity *in-vivo.*

In addition, export of mRNAs from the nucleus involves a quality-control mechanism that checks if mRNAs are fully spliced and have properly formed 5’ and 3’ ends^24^. I hypothesized that mRNA export may involve a quality control that removes mRNAs with errors. I used the ENCODE dataset in which nuclear and cytoplasmic poly-A+ mRNAs I re sequenced, thus I can compare nuclear and cytoplasmic fractions from the same cell line grown in the same conditions and processed in the same manner. I find that the nuclear fraction has a higher RNA polymerase error rate than does the cytoplasmic fraction **(Figure 2c,d)**, suggesting that either that nuclear RNA-seq has a higher technical error rate or that the cell has mechanisms for reducing the effective polymerase error rate by preventing the export of mRNAs that contain errors.

Rpb9 and Dst1 are known to be involved in RNA polymerase fidelity *in-vitro,* yet there is conflicting evidence as to the role of Dst1 *in-vivo*^6,15,23,25-27^. Part of these conflicts may result from the fact that the only available assays for RNA polymerase fidelity are special reporter strains that rely on DNA sequences known to increase the frequency of RNA polymerase errors. While I found that RPB9 expression correlates with RNA-seq error rates in mammalian cells, correlation is not causation. Furthermore, differences in RNA levels do not necessitate differences in stoichiometry among the subunits in active Pol II complexes. In order to determine if differential expression of RPB9 or DST1 are causative for differences in RNA polymerase fidelity in-vivo, I constructed two yeast strains in which I can alter the expression of either RPB9 or DST1 using B-estradiol and a synthetic transcription factor that has no effect on growth rate or the expression of any other genes^28,29^. I grew these two strains (Z_3_EV_pr_-RPB9 and Z_3_EV_pr_-DST1) in different concentrations of B-estradiol and performed RNA-seq. I find that cells expressing low levels of RPB9 have high RNA polymerase error rates **(Figure 3a)**. Likewise, cells with low DST1 have high error rates **(Figure 3a)**. The increase in errors rate is not a property of cells defective for transcription elongation **(Figure 3 - figure supplement 1)**. The increase in error rates due to mutations in Rpb9 and Dst1 have not been robustly measured, however, there are some rough numbers. Here, the measured increase in error rate is 13%, while the measured effect of Rpb9 deletion *in-*vitro is 5-fold^38^ and in-vivo following reverse transcription is 30%^25^. If 2% of the observed mismatches are due to RNA polymerase errors, a 5-fold increase in polymerase error rate results in a 10% increase in measured mismatch frequency; this is consistent with RNA polymerase fidelity of 10^−6^-10^−5^ and overall RNA-seq error rates of 10^−4^. Note that in our assay cells still express low levels or RPB9, and we therefore expect the increase in error rate to be lower, suggesting that RNA polymerase errors constitute 5-10% of the measured mismatches. Our ability to genetically control the expression of DST1 and RPB9, and measure changes in RNA-seq error rates is consistent with MORPhEUS measuring RNA polymerase fidelity. In addition, genetic reduction in RNA polymerase fidelity results in increased intron retention, consistent with RNA polymerase errors causing reduced splicing efficiency **(Figure 3b)**.

**Figure 3.**
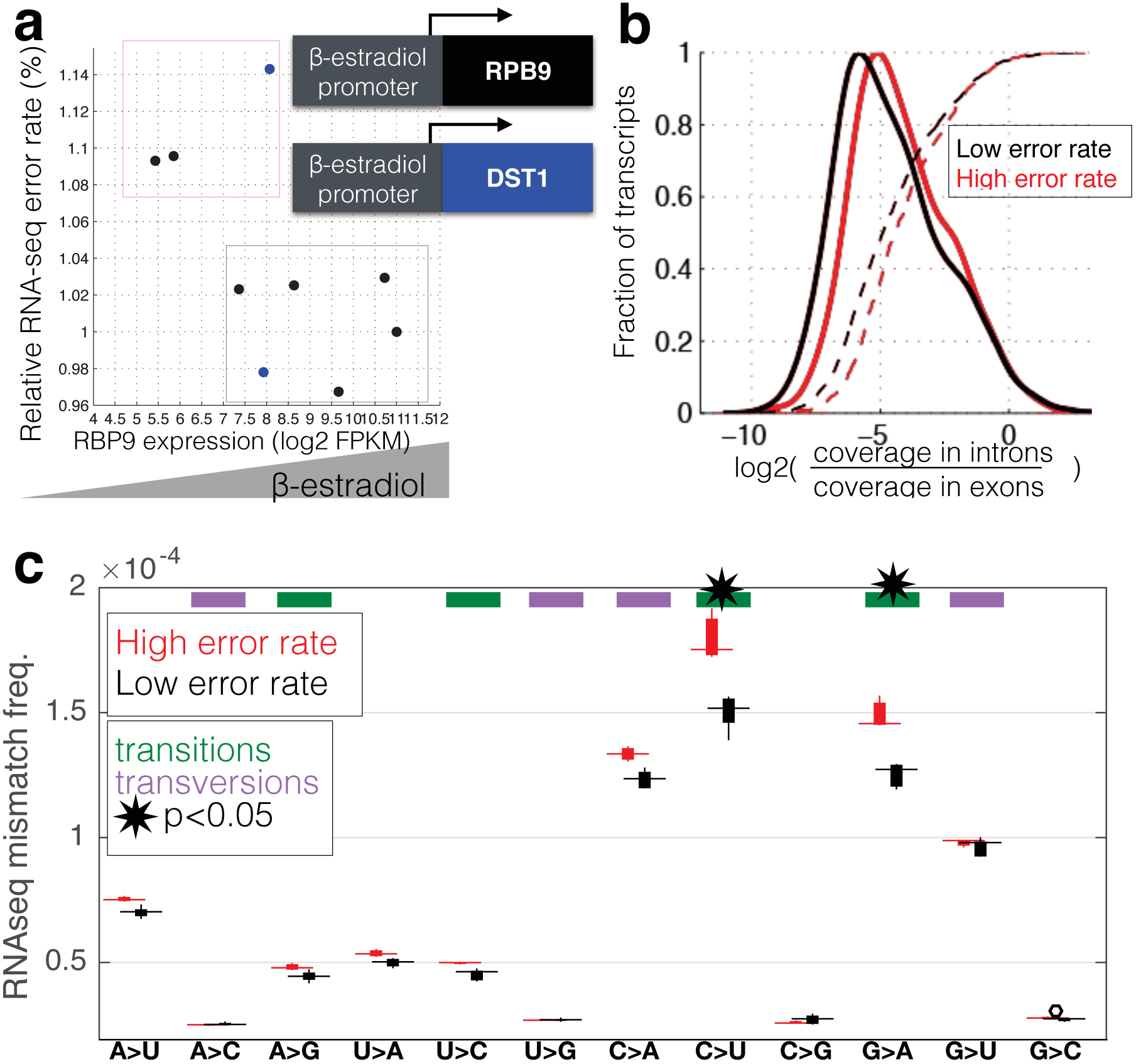
RNA polymerase error rate is determined by the expression level of RPB9 and DST1. **(a)** RNA-seq error rates I re measured for two strains (Z_3_EVpr-RPB9, black points, Z_3_EVpr-DST1, blue points) grown at different concentrations of β-estradiol. The points show the relationship between RPB9 expression levels (determined by RNA-seq) and RNA-seq error rates. The blue points show RPB9 expression levels for the Z_3_EVpr-DST1 strain, in which DST1 expression ranges from 16 FPKM at 0nM β-estradiol to 120 FPKM native expression to 756 FPKM at 25nM β-estradiol. Low induction of both DST1 or RPB9 results in high RNA-seq error rates (red box), while wild-type and higher induction levels result low RNA-seq error rates (black box). **(b)** Across all genes, the intron retention rate is higher in conditions with low RNA polymerase fidelity (t-test between high and low error rate samples, p=0.029), consistent with the hypothesis that RNA polymerase errors result in splicing defects. **(c)** The error rate for each of the 12 single base changes are shown for induction experiments that gave high (red) or low (black) RNA-seq error rates. Transitions (G<->A, C<->U) are marked with green boxes and transversions (A<->C, G<->U) with purple

A unique advantage of MORPhEUS is that it measures thousands of RNA polymerase errors across the entire transcriptome in a single experiment, and thus enables a complete characterization of the mutation spectrum and biases of RNA polymerase. I asked how altered RPB9 and DST1 expression levels affect each type of single nucleotide change. I find that, with decreasing polymerase fidelity, transitions increase more than transversions, and that C->U errors are the most common **(Figure 3c)**. This result, along with other sequencing based results^4^, have shown that DNA and RNA polymerase have broadly similar error profiles^2^; it will be interesting to see if all polymerases share the same mutation spectra, and if this is due to deamination of the template base, or is a structural property of the polymerase itself. Interestingly, I find that coding sequences have evolved so that errors are less likely to produce in-frame stop codons than out-of-frame stop codons, suggesting that natural selection may act to minimize the effect of polymerase errors **(Figure 4)**.

**Figure 4.**
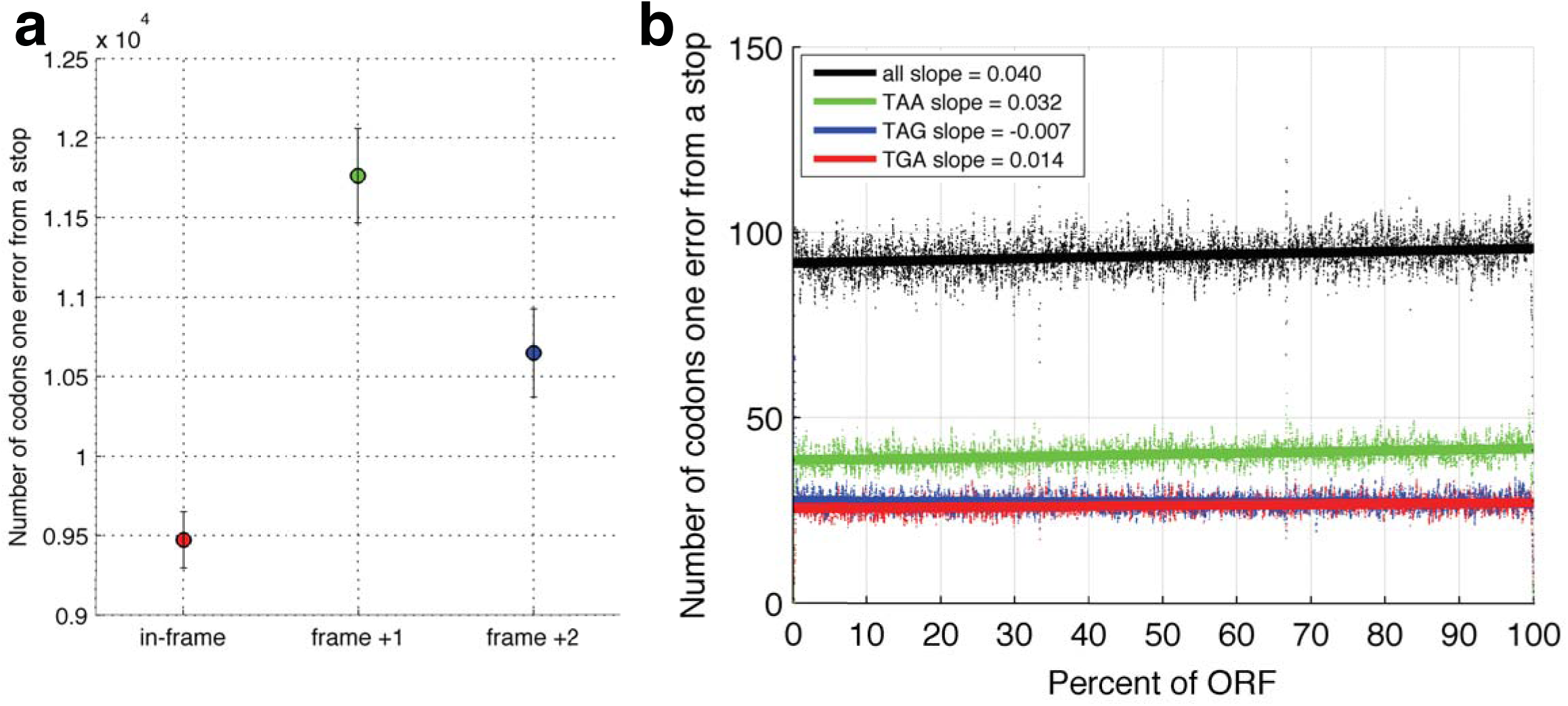
In-frame stop codons are less likely to be created by polymerase errors. For all genes in yeast, I calculated the number of codons which are one polymerase error from a stop codon. **(a)** Fewer in-frame codons can be turned into a stop codon by a single nucleotide change, compared to out-of-frame codons. **(b)** Codons that are one error away from generating an in-frame stop codon are more likely to be found at the ends of ORFs, compared to the beginning of the ORF.

Here I have presented proof that relative changes in RNA polymerase error rates can be measured using standard Illumina RNA-seq data. Consistent with previous work *in-vivo* and *in-vitro,* I find that depletion of RPB9 or Dst1 results in higher RNA polymerase error rates. Futhermore, I find that expression of RPB9 negatively correlates with RNA-seq error rates in human cell lines, suggesting that differential expression of RPB9 may regulate RNA polymerase fidelity in-vivo in humans. In addition, consistent with the errors detected by MORPhEUS being due to RNA polymerase and not technical errors, in reads spanning an exon-intron junction, the measured error rate is higher at the 5’ donor splice site, suggesting that RNA polymerase errors result in intron retention. Because it can be run on existing RNA-seq data, I expect MORPhEUS to enable many future discoveries regarding both the molecular determinants of RNA polymerase error rates, and the relationship between RNA polymerase fidelity and phenotype.

## Acknowledgements

I thank members of the Carey lab and the computational genomics groups in the PRBB for thoughtful discussions.

## Materials and methods

### Counting RNA polymerase errors in already aligned ENCODE data

Much existing RNA-seq data is available as bam files aligned to the human genome. In order to bypass the most computationally expensive step of the pipeline, I developed a method capable of using RNA-seq reads aligned with spliced aligners. First, in order to avoid increased mismatch rates at splice junctions due to alignment problems with both spliced and unspliced reads, I used samtools^30^ and awk to remove all alignments that don’t align along the full length of the genome (eg: for 76bp reads, only reads with a CIGAR flag of 76M). The remaining reads I re trimmed (bamUtil, trimBam) to convert the first and last 10bp of each read to Ns and set the quality strings to ‘!’. I then used samtools mpileup (-q30 –C50 –Q30) and custom perl code to count the number of reads and number of errors at each position in genome. Positions with too many errors (eg: more than one read of the same non-reference base) I re not counted.

### Measurement of error rates at splice junctions

I used the UCSC table browser^31^ to download two bed files: hg19 EnsemblGenes introns with -10bp flanking from each side, and another file with the introns and +10bp flanking on either side. I then used bedtools^32^ (bedtools flank -b 20 -l 0 & bedtools flank -l 20 -b 0) to generate bed files with intervals that contain the splicing donor and acceptor sites, respectively. In addition, I used bedtools getfasta on the +10bp flanking bed file to keep only introns flanked by GT and AG donor and acceptor sites. The final result is a pair of bam files with intervals centered on the splicing donor or acceptor sites. I used this new bed file to count error rates around each splice junction. The error rate at each position (eg: -10, -9, - 8, etc from the G at the 5’ donor site) is the sum of all errors at that position, divided by the sum of all reads. Positions are relative to the splicing feature, not to the genome, as error rates at any single genomic position are dominated by sampling bias. Per mono, di and tri-nucleotide background error rates I re calculated using the same scripts, but without limiting mpileup to the splice junctions.

### Strain construction and RNA sequencing for RPB9 and DST1 strains

The parental strain DBY12394^33^ (GAL2+s288c repaired HAP1, ura3Δ, leu2Δ0::ACT1pr-Z3EV-NatMX) was transformed with a PCR product (KanMX-Z3EVpr) to generate a genomically integrated inducible RPB9 (LCY143) or DST1 (LCY142). Correct transformants I re confirmed by PCR. To induce various levels of expression, strains I re grown in YPD + 0,3,6,12 or 25nM β-estradiol (Sigma E4389) for more than 12 hours to a final OD_600_ of 0.1 – 0.4. Cellular RNA was extracted using the Epicenter MasterPure RNA Purification Kit, and Illumina sequencing libraries I re prepared using the Truseq Stranded mRNA kit, and sequenced on a HiSeq2000 with at least 20,000,000 50bp sequencing reads per sample.

I used bwa^34^ (-n 2, to permit no more than two mismatches in a read) to align the yeast RNA-seq reads to the reference genome, and trimBam from bamUtil to mask the first and last 10bp of each read. I used samtools mpileup^30^ (-q 30 -d 100000 - C50 –Q39) to count the number of reads and mismatches at each position in the genome, discarding low confidence mapping, reads that map to multiple positions, and low quality reads. Duplicate reads can be removed from the fastq file if the coverage is low enough so that all unique read sequences are expected to come the same RNA fragment; this is the case for low coverage paired-end reads with a variable insert size, but not for very high coverage datasets or single-ended reads.

### pre-existing RNA-seq datasets

For the intron retention analysis in human cells, data are from NCBI SRA PRJNA253670. Data for the elc4 and spt4 analysis are from PRJNA167772 and PRJNA148851, respectively. For RPB9 correlation, ENCODE35 data (SRA PRJNA30709) are all from the Gingeras lab at CSHL.

---

Figures

**Figure 1 – figure supplement 1.**
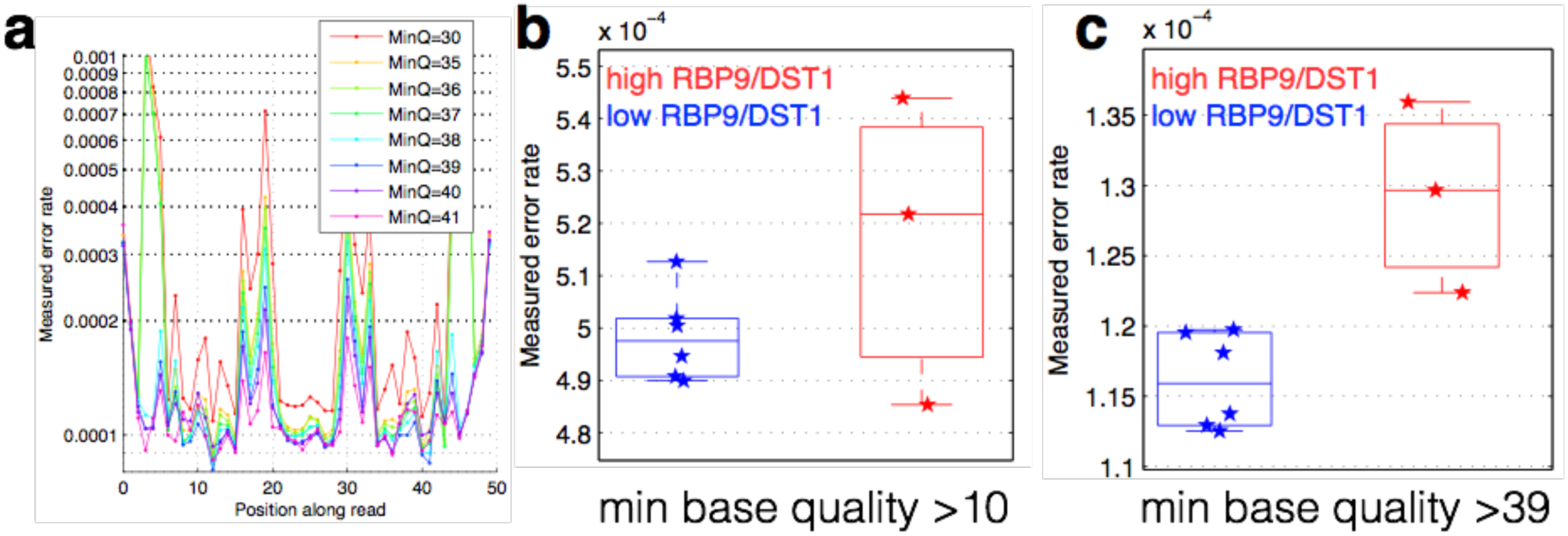
Cycle-specific error rates and better differentiation of genetically determined error rates using base quality value cutoffs. Six yeast RNA-cDNA libraries were sequenced on the same lane in a HiSeq. **(a)** The average mismatch rate (across the six cDNA libraries) to the reference genome at each position was determined using different minimum base-quality thresholds using GATK ErrorRatePerCycle. Independent of the quality threshold, cycles at the ends, as well as some cycles in the middle, have high error rates. **(b)** The measured error rate for each sample using a minimum base quality of 10. **(c)** The measured error rate for each sample using a minimum base quality of 39.

**Figure 1 – figure supplement 2.**
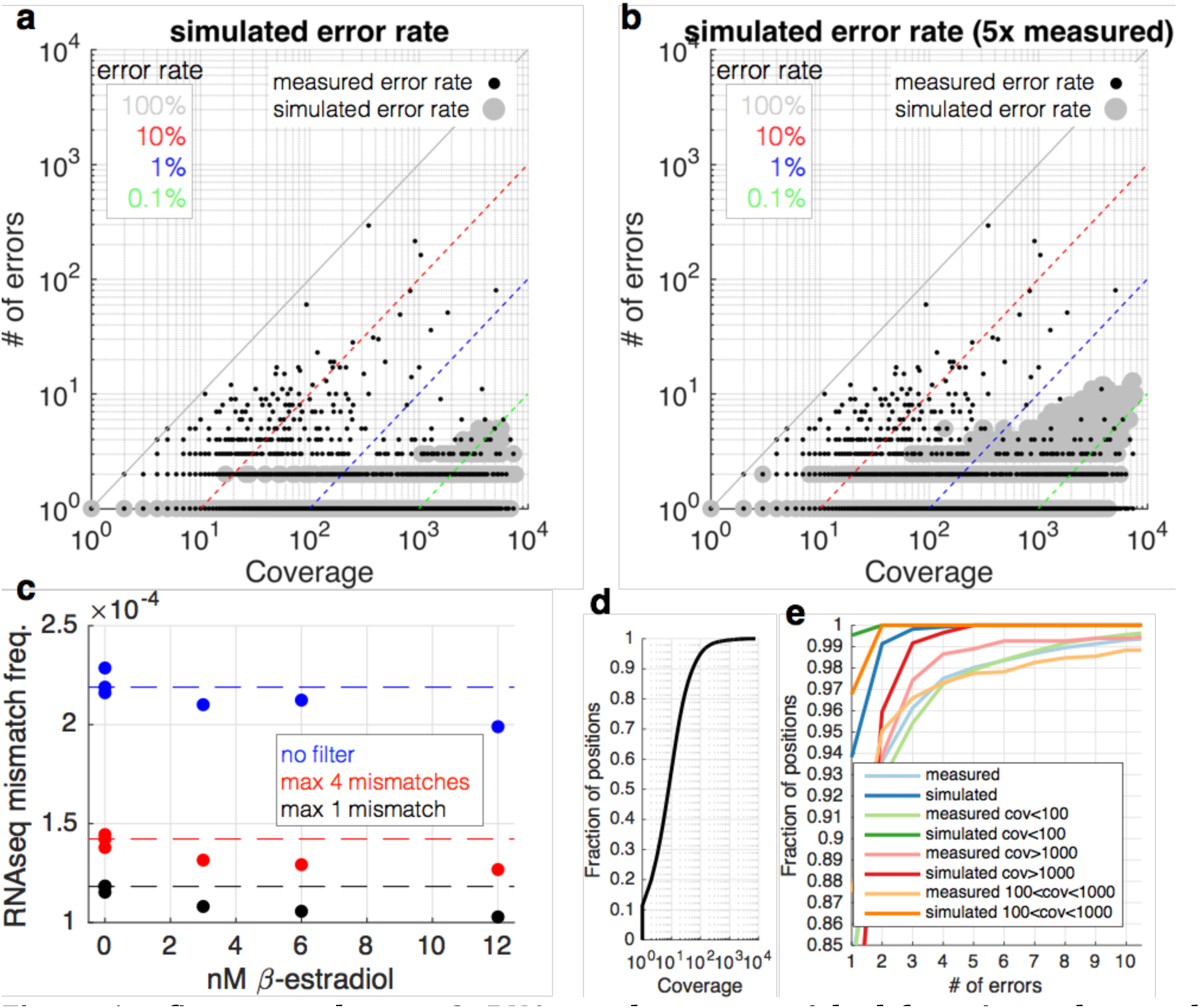
RNA-seq data are enriched for mismatches to the reference genome that occur far more often than expected. **(a)** At each coverage (x-axis), a point is shown if there is any positions in the genome with the observed number of errors (y-axis) (small black dots). The diagonal lines show mismatch frequencies of 100%, 10%, 1% and 0.1% — any point falling on these lines has the given mismatch frequency. With large grey circles are shown simulated data in which the same coverage as the yeast RNA-seq data are used, but with a mismatch frequency identical to the measured overall mismatch frequency of the yeast data. Locations in the graph in which a black point occurs but there is no grey point are locations in which there are more mismatches than expected by change. Note that at a coverage of less than 100, we expect to see no mismatches more than twice, and 0.5% of positions with 2 observances of identical mismatches. **(b)** Identical to (a) but with the simulated mismatch frequency 5x the observed. **(c)** Shown are measured mismatch frequencies for the yeast RPB9 and DST1 induction data at different B-estradiol concentrations, at different filters for the maximal allowed number of observed identical mismatches. The dashed lines show the average mismatch frequency for the 0nM condition. For all filters, low B-estradiol conditions have higher RNA-seq mismatch frequencies. **(d)** The coverage of the yeast RNA-seq data; ~95% of the genome is covered by less than 100 reads. **(e)** Shown are the fraction of positions in the genome (y-axis) with X errors (x-axis) for the yeast RNAseq data (cyan) and simulated data (blue). Also shown are the same data for positions of the genome with different coverage. For positions covered by less than 1000 reads (95% of the genome) the expectation is 0 or 1 sequence mismatch (blue and orange lines). Positions with far greater numbers of mismatches are likely due to sub-clonal mutations and technical bias.

**Figure 2 – figure supplement 1.**
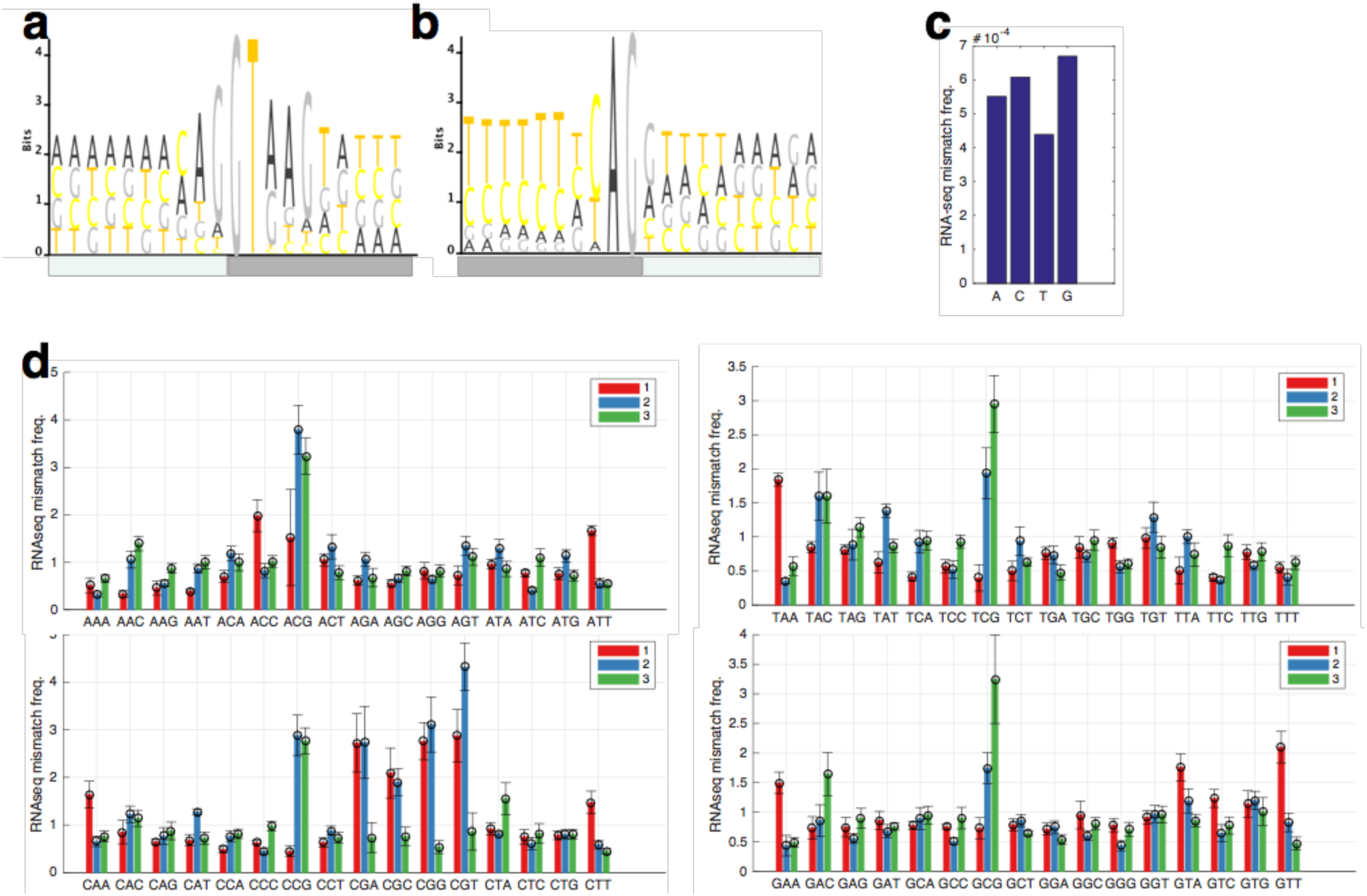
RNA-seq mismatch rates for all trinuculeotides in chromatin associated and nuclear RNAs. **(a,b)** The 5’ and 3 splicing motifs in the human genome. **(c)** The RNA-seq mismatch frequencies for all single nucleotides. **(d)** The RNA-seq mismatch rate to the reference genome for each trinucleotide, normalized to the average mismatch rate across all trinucleotides. For each trinucleotide, red shows the mismatch frequency at the first base, blue at the second, and green at the third. Error bars are standard deviation across all samples.

**Figure 2 – figure supplement 2.**
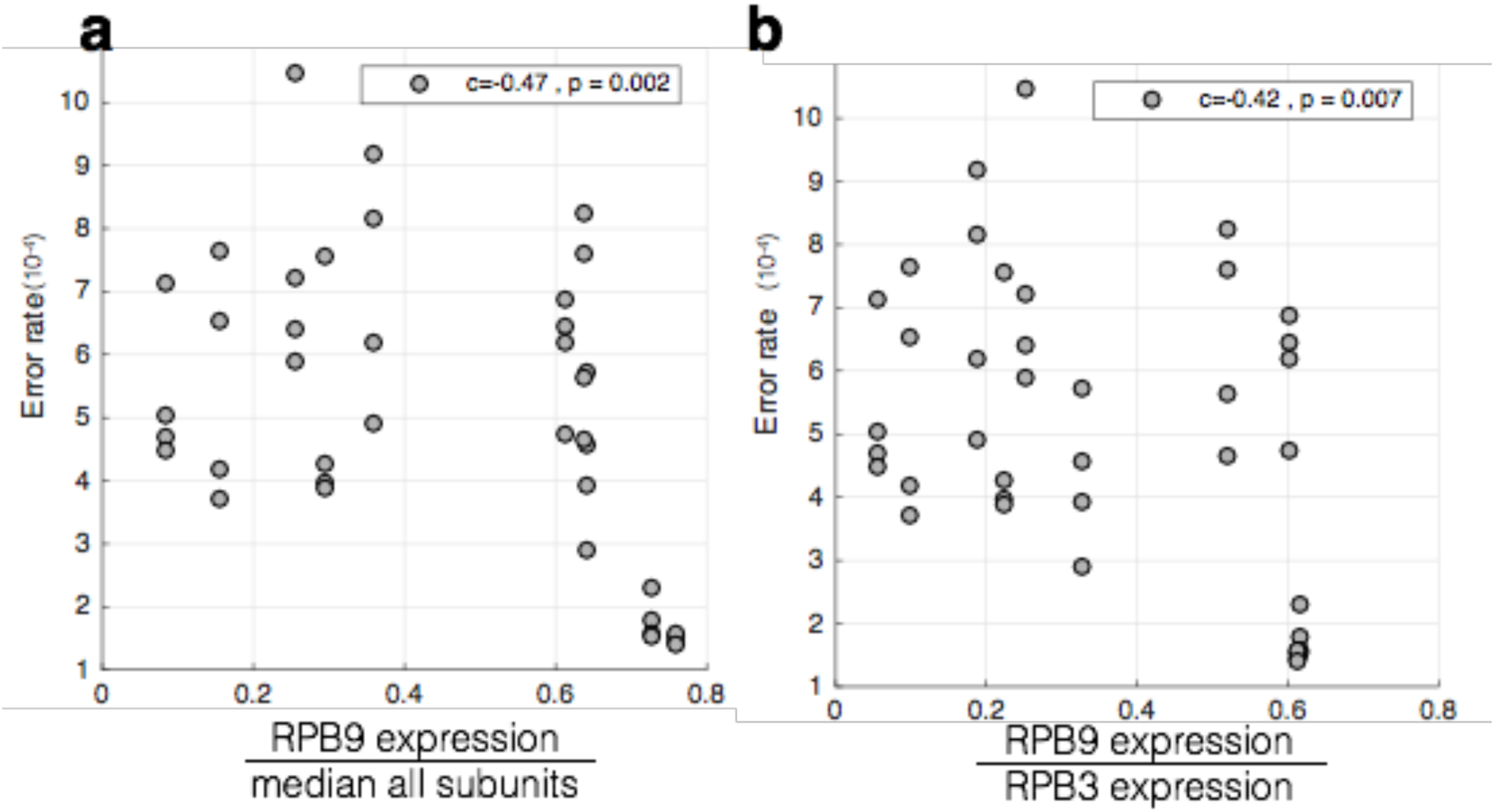
RBP9 expression negatively correlates with RNA-seq mismatch rates. The mismatch frequency is shown across all cells lines. **(a)** RPB9 mRNA expression is normalized by the median expression level of all subunits. **(b)** RPB9 mRNA expression is normalized by RBP3 (POLR2C) expression.

**Figure 3 – figure supplement 1.** Decreases in RPB9 and DST1 expression in yeast results in more single base insertions in RNA-seq data. For each RNA-seq dataset, the number of inserts (+N) or deletions (−N) in the mpileup output (N is the number o bases in the indel) were counted, and this number divided by the total number of mapped reads in each sample. On the right are the same data but zoomed in on each metric to better show the comparison between the two sets of samples.

**Figure 3 – figure supplement 2.**
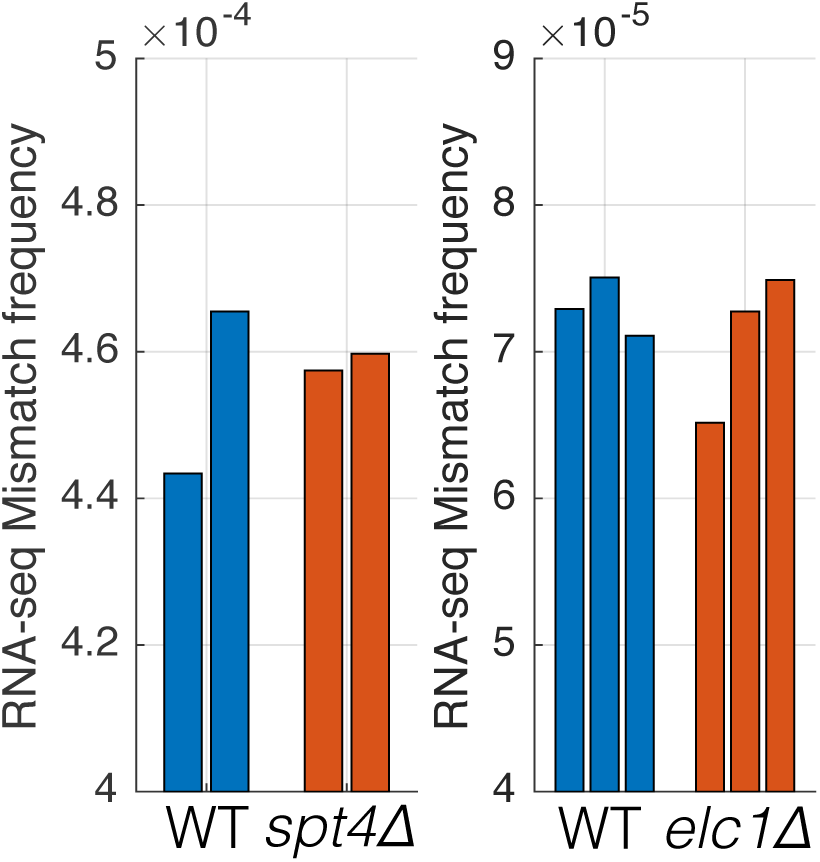
Mutations that affect transcription elongation do not affect measured RNA-seq mismatch frequencies. Two separate experiments were performed with wild-type controls and mutants involved in transcription elongation. Individual bars show the RNA-seq mismatch frequency of biological replicates.

